# Dopamine-Modified Hyaluronic Acid (DA-HA) As A Novel Dopamine-Mimetics With Minimal Autoxidation And Cytotoxicity

**DOI:** 10.1101/2022.08.21.504712

**Authors:** Sunpil Kim, Ye-Ji Kim, Kyoung Hwan Park, Kang Moo Huh, Sun-Woong Kang, C. Justin Lee, Dong Ho Woo

## Abstract

Dopamine-modified hyaluronic acid (DA-HA) has been initially developed as an efficient coating and adhesion material for industrial uses. However, the biological activity and safety of DA-HA in the brain have not been explored yet. Here, we report a series of evidence that DA-HA exhibits similar functionality as dopamine (DA), but with much lower toxicity arising from autoxidation. DA-HA shows very little autoxidation even after 48-hour incubation. This is profoundly different from DA and its derivatives including L-DOPA, which all induce severe neuronal death after pre-autoxidation, indicating that autoxidation is the cause of neuronal death. Furthermore, *in vivo* injection of DA-HA induces significantly lower toxicity compared to 6-OHDA, a well-known oxidized and toxic form of DA, and alleviates the apomorphine-induced rotational behavior in the 6-OHDA animal model of Parkinson’s disease. Our study proposes that DA-HA with DA-like functionalities and minimal toxicity can be an effective therapeutic substitute for L-DOPA in Parkinson’s disease.

## Introduction

Dopamine (DA) is one of the major neurotransmitters in the brain. Dopaminergic signaling is involved in various physiological functions such as movement ^1^, reward ^2^, learning ^3^, and memory ^4,5^. The nigrostriatal pathway is known as the major dopaminergic signaling pathway implicated in voluntary movement control and Parkinson’s disease with degeneration of dopaminergic neurons in this pathway ^6^. To treat this debilitating disease, clinicians have tried to supply DA into the brain. Unfortunately, DA cannot penetrate the blood-brain barrier (BBB). Therefore, there has been a pressing need to develop other ways to deliver DA or DA-like drugs into the brain.

The most widely used drug to treat Parkinson’s disease is levodopa (L-DOPA), which is a precursor metabolite of DA and has an excellent BBB-permeability ^7^. L-DOPA improves motor symptoms at the initial stages of disease progression. However, it has been well-known that long-term treatment of L-DOPA precipitates severe side effects called L-DOPA-induced dyskinesia (LID) ^8^, implying that L-DOPA itself has serious toxicity. It has been proposed that the toxicity of DA and its derivatives is primarily caused by either intracellular free radicals of reactive oxygen species (ROS) ^9^ or extracellular oxidation of DA ^10^ or both. On one hand, when DA enters the cell inside via monoamine transporters, DA is degraded by monoamine oxidases and induces cytotoxicity by a formation of radicals through unpaired electrons of generated superoxide ^11^. On the other hand, DA is oxidized to generate external free radicals through extracellular oxidation and causes cytotoxicity on diverse cell types such as mesencephalic neurons ^12,13^, cortical neurons ^14^, and astrocytes ^15^, implying that external oxidation of DA induces non-specific cell death. However, it still remains unclear which of the two proposed mechanisms, intracellular or extracellular ROS, is the major cause of DA-induced cytotoxicity.

External free radicals are generated through a process called autoxidation, a self-oxidation without cells in an aqueous solution ^16^. Autoxidation of DA can be easily measured by optical density (OD) at wavelengths ranging from 350 to 450 nm ^15^. Interestingly, it has been reported that DA-induced cortical neuronal cell death is accelerated with nomifensine, a monoamine transporter inhibitor ^14^. Moreover, both an inhibitor for superoxide dismutase and deferoxamine mesylate, an iron chelator, are shown to block DA-induced cell death ^14^. These reports strongly suggest that autoxidation of DA induces neuronal cell death. Unlike DA, DA-modified hyaluronic acid (DA-HA) has been proven to be safe in biological applications. DA-HA was originally developed as an adhesive for attaching electrodes to the surface because of its high bio-compatibility ^17,18,19^. DA-HA is synthesized by a polymer reaction using EDC / NHS (1-ethyl-3-(3-dimethylaminopropyl) carbodiimide / N-hydroxysuccinimide) to conjugate DA to the carboxyl residue of hyaluronic acid (HA) polymer ^20^. The main benefit of employing HA is that it slows down the rate of degradation ^21^. DA-HA conjugation has been also utilized to stably release other drugs for a long period of time with high bio-compatibility ^22^. However, the physiological role and mechanism of DA-HA action in the brain have not been studied. Furthermore, autoxidation of DA-HA has not been investigated yet.

In this study, we firstly investigate the functionality and autoxidation of DA-HA and its potential application in the brain, especially in the animal model of Parkinson’s disease. Using GRAB^_DA2m_^ imaging, *ex vivo* Ca^2+^ imaging, cell viability test, autoxidation measurement, *in vivo* DA neuron toxicity, and apomorphine-induced rotation test in 6-OHDA lesion mouse, we demonstrate that DA-HA shows minimal autoxidation with virtually no cytotoxicity. Furthermore, supplementation of DA-HA rescues a motor symptom of the 6-OHDA animal model of Parkinson’s disease. Based on these findings, we propose DA-HA as a potential therapeutic substitute for L-DOPA to treat patients suffering from Parkinson’s disease.

## Results

### Synthesis and validation of DA-HA

The conjugation method of heparin ^23^ and HA ^24^ has been well established and this conjugation can be validated by nuclear magnetic resonance (NMR), scanning electron microscopy (SEM), and atomic force microscopy (AFM). DA was conjugated with the carboxyl group of hyaluronic acid (HA)(Fig. 1a and Supplementary Fig. 1) as previously described ^20^. The structure of the DA-HA conjugate was characterized by using ^1^H-NMR (Supplementary Fig. 1b, c). The protons of the aromatic ring of DA showed peaks at 6.6-6.9 ppm (Peak 5 of DA-HA conjugate in structure and spectrum, dot box in Supplementary Fig. 1d) and those of N-acetyl groups of HA showed 1.9 ppm of the spectrum of ^1^H-NMR (Peak 1 of DA-HA conjugate in structure and spectrum, Supplementary Fig. 1d), confirming an appropriate conjugation of DA to HA. Using Fourier transform infrared spectroscopy-attenuated total reflectance (FTIR-ATR)^25^, we measured peak changes of 1720, 1630, and 1410 cm^-1^ which indicated a DA conjugation (3 arrows in the blue trace, Supplementary Fig. 1c). As a result, 5.33, 23.59, 35.33, and 39.02 mg of DA amount exists in 1 g HA, demonstrating 0.1, 0.2, 0.5, and 1 of feeding DA to HA molar ratio, respectively (Supplementary Fig. 1e and Table 1). In addition, the oxidation processes and polymerization of DA and its derivatives used in this study are schematized (Fig. 1b-d). Taken together, these results validated that we successfully synthesized DA-HA.

**Fig. 1.**
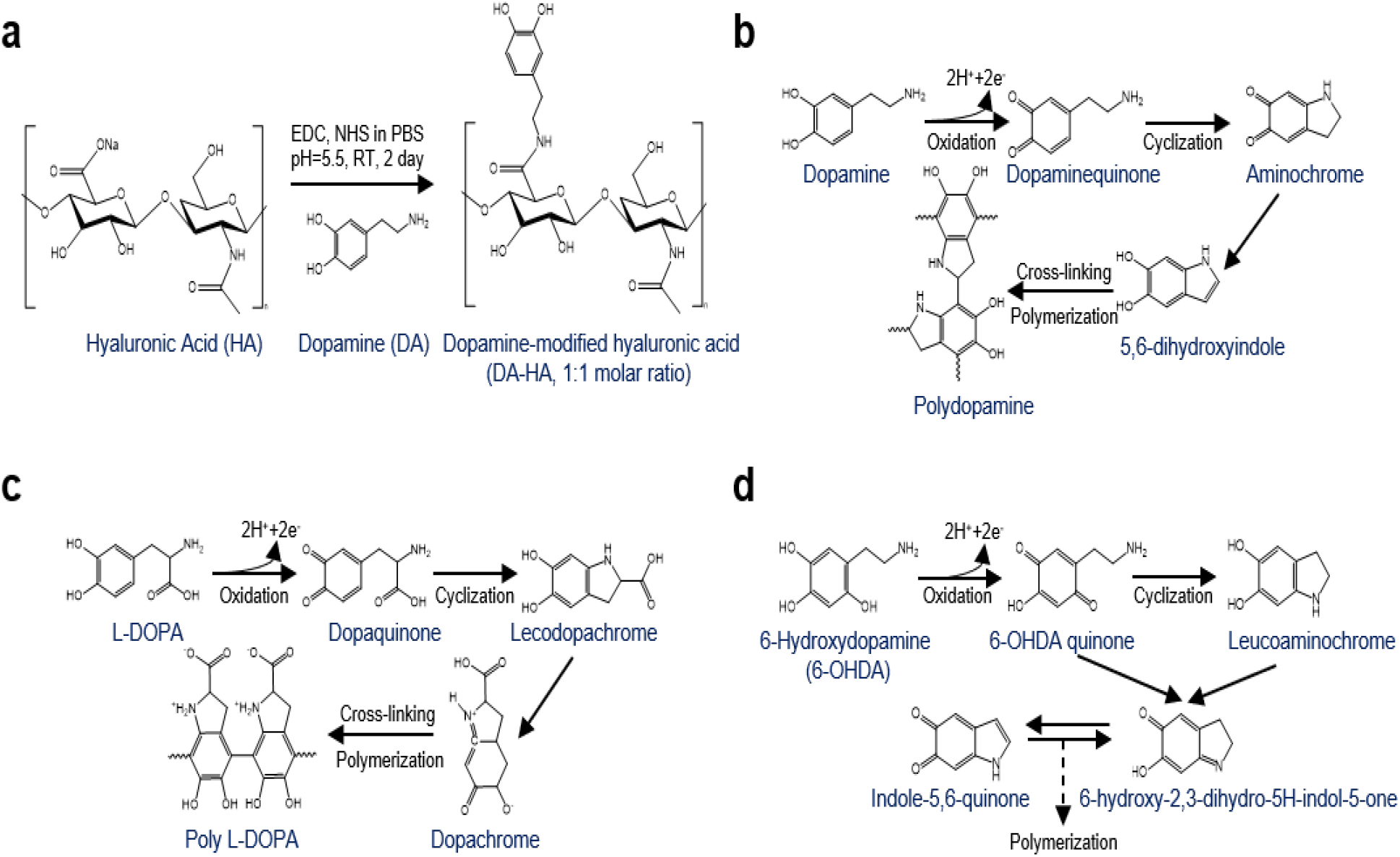
Process of DA-HA synthesis and oxidation and polymerization processes of DA and its derivatives. **a** A schematic diagram of the process of DA-HA synthesis. **b-d** Oxidation and polymerization processes of DA (**b**), L-DOPA (**c**), and 6-OHDA (**d**).

### Functional characterization of DA-HA and DA

To investigate whether DA-HA shows similar pharmacological properties as DA, we utilized a recently developed DA sensor, GRAB_DA2m_ ^26^. We transfected AAV-GFAP104-GRAB_DA2m_ clone as an exogenous DA receptor into rat primary cultured astrocytes and perform real-time fluorescence imaging (Fig. 2a). DA content of DA-HA was calculated by DA concentration. Fluorescence transients were gradually increased by bath application of either DA or DA-HA in a dose-dependent manner (Fig. 2b, c). Half maximal effective concentration (EC_50_) of DA and DA-HA were 0.077 µM and 0.062 µM, respectively (Fig. 1d), indicating similar EC_50_ values. In addition, we observed similar changes in GRAB_DA2m_ transients by the same concentration of DA and DA-HA onto the same cell (Fig. 2e, f). These results indicated that DA-HA showed similar functionality as DA *in vitro*. To recapitulate these results *ex vivo*, we injected AAV-GFAP104-GRAB_DA2m_ virus into nucleus accumbens (NAc) including core (AcbC) and shell (Acbsh) regions, and performed a fluorescence imaging experiment (Fig. 2g). Both 10 µM DA and DA-HA, but not HA, induced similar peak amplitude of fluorescence transients from the same region of interest (ROI, Fig. 2h, i). These results indicated that DA-HA shows similar pharmacological properties as DA *ex vivo*.

**Fig. 2.**
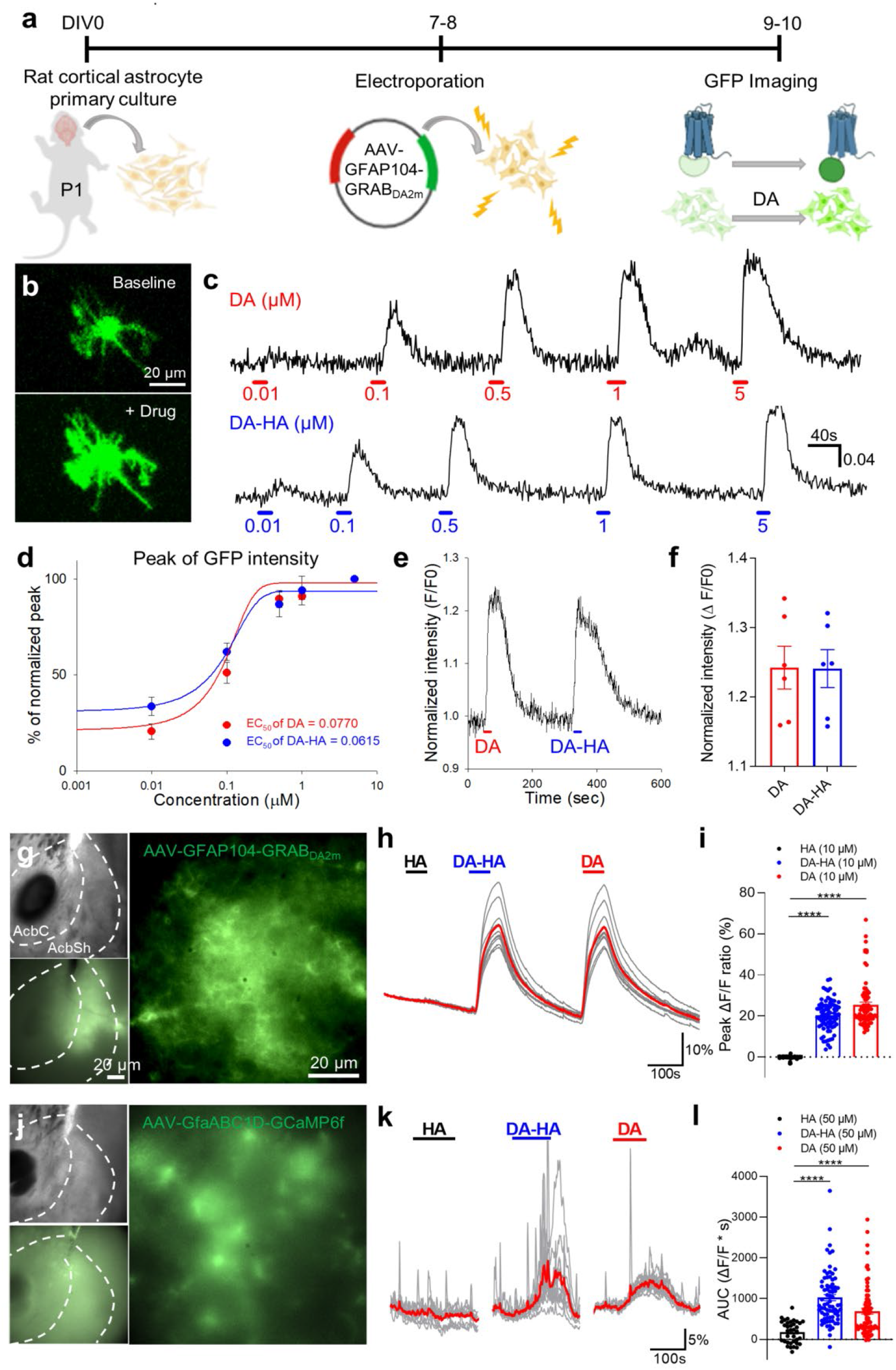
Functional characterization of DA-HA and DA by GRAB_DA2m_ DA sensor and GCaMP6f Ca^2+^ sensor. **a** A schematic diagram for an experimental schedule of GFP imaging in AAV-GFAP104-GRAB^DA2m^ transfected astrocyte. **b** Intensity of GRAB^DA2m^ baseline (top) and that after adding DA (bottom). Scale bar: 20 µm. **c** Representative traces of GFP transients with a dose of 0.01, 0.1, 0.5, 1, and 5 µM of DA (top) and DA-HA (bottom) from AAV-GFAP104-GRAB^DA2m^ transfected astrocytes. **d** Dose-dependent curves for the normalized peak of GFP transients and EC^50^ by DA (red, n = 3) and DA-HA (blue, n = 4). **e** Normalized GRAB^DA2m^ transient of 1 µM DA and 1 µM DA-HA in the same astrocyte. **f** Summary bar graph for normalized peak amplitude from **e** (n = 6 for each condition). Paired t-test, p = 0.83. **g** Expression of AAV-GFAP104-GRAB^DA2m^ virus in NAc. Dotted lines indicate accumbens core (AcbC, Inside) and accumbens shell (AcbSh). Scale bar: 200 µm and 50 µm. **h** Representative traces of GRAB^DA2m^ under treatment of 10 µM HA, DA-HA, and DA. Red trace indicates averaged trace. **i** Summary bar graph of the peak amplitude of GRAB^DA2m^ transients from **h** (n = 79 ROIs for each condition). One-way ANOVA, F(2, 234) = 228.4, p < 0.0001; HA vs DA-HA and HA vs DA, ****p < 0.0001. **j** Expression of AAV-GfaABC1D-GCaMP6f virus in NAc. **k** Representative traces of Ca^2+^ transients under treatment of 50 µM HA, DA-HA, and DA. Red traces indicate averaged traces. **l** Summary bar graph of the area under curve (AUC) analysis of Ca^2+^ transients from **k** (n = 36, 75, and 75 ROIs). One-way ANOVA, F(2, 183) = 26.12, p < 0.0001; HA vs DA-HA and HA vs DA, ****p < 0.0001. Tukey’s multiple comparisons test for **i** and **l**.

It has been previously demonstrated that astrocytes in NAc show Ca^2+^ transients upon synaptic DA release ^27^. To investigate whether DA-HA can evoke DA-induced Ca^2+^ transient in NAc astrocytes, we injected AAV-GfaABC1D-GCaMP6f virus into NAc and monitor NAc astrocytic Ca^2+^. Both 50 µM DA and DA-HA, but not HA, induced similar Ca^2+^ transients from the same ROI (Fig. 2k, l), indicating that DA-HA functionally activates endogenous DA receptor in NAc astrocyte. In conclusion, DA-HA can activate exogenous GRAB_DA2m_ sensor as well as endogenous DA receptor in NAc astrocyte.

### Long-term stability and safety of DA-HA on cultured neurons

It has been previously reported that 10 to 100 µM DA induces rat cortical neuronal cell death via the mechanism of forming ROS ^14^. To investigate whether DA-HA induces cell death or not, we incubated 10 µM DA, 10 µM DA-HA, and 120 µM HA for 7 days to measure cell viability (Fig. 3a). By counting the number of NeuN^+^ cells, we observed that DA significantly decreased the number of NeuN^+^ cell on day 7 but not day 2 (Fig. 3b, c). In contrast, DA-HA, as well as HA, did not induce any significant neuronal death on day 7 (Fig. 3b, c). To see if the conjugation of DA to HA is critical for the protective effect, we incubated cultured neurons with unconjugated HA and DA together (DA + HA). We found that the unconjugated HA and DA treated group showed a similar level of toxicity as DA treated group (Fig. 3c), indicating that the conjugation of DA-HA is important for the protective effect of DA-HA.

**Fig. 3.**
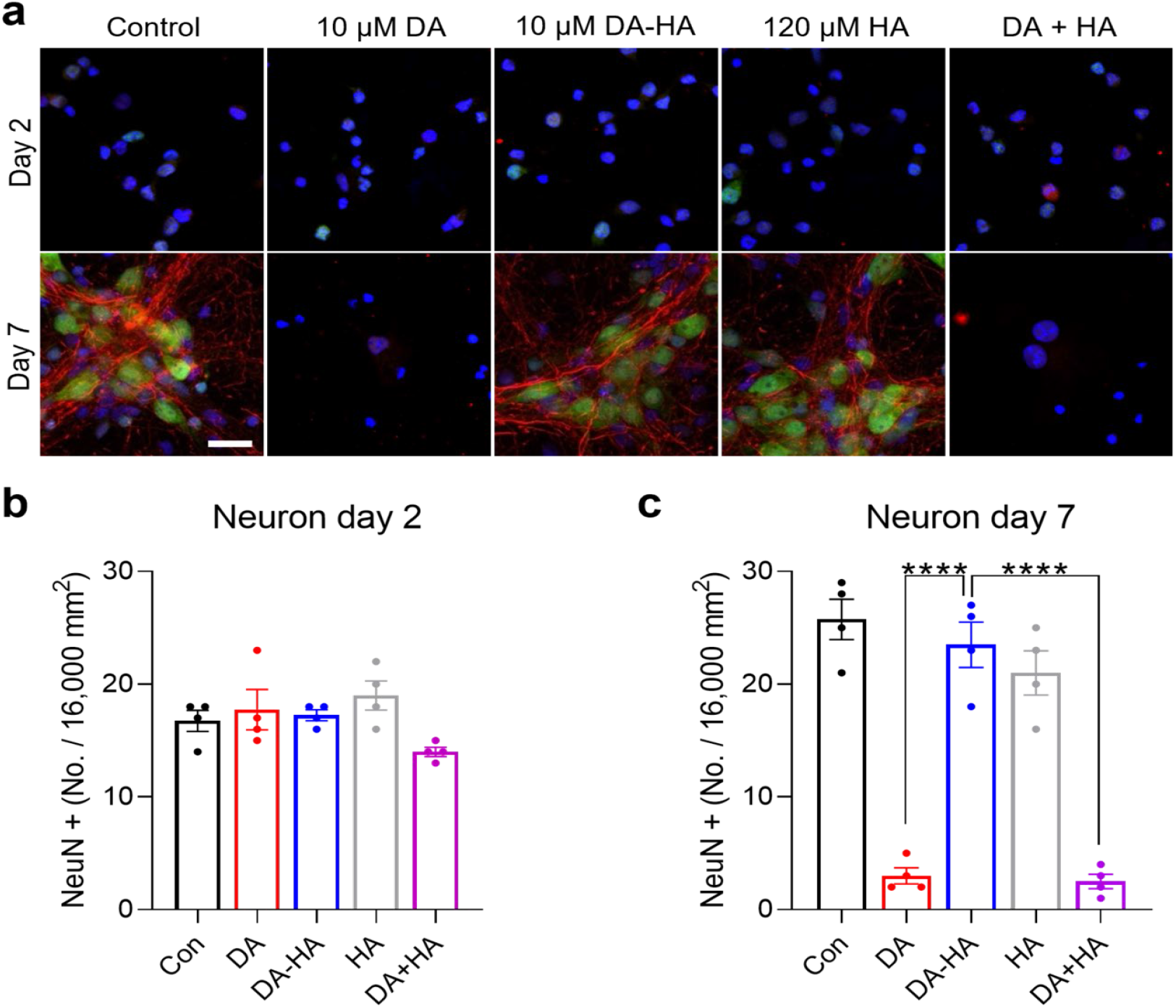
Conjugation of DA and HA is critical for safety. **a** Merged immunostaining images of cultured neurons with NeuN (green), neurofilament (NF, red), and DAPI (blue) with no treatment, 10 µM DA, 10 µM DA-HA, 120 µM HA, and 10 µM DA + 120 µM HA at day 2 and day 7. **b, c** Summary bar graph of NeuN+ cells at day 2 (**b**, n = 4 for each condition) and day 7 (**c**, n = 4 for each condition). One-way ANOVA, F(4, 15) = 54.3, p < 0.0001; DA-HA vs DA and DA-HA vs DA+HA, ****p < 0.0001 in (**c**). Dunnett’s multiple comparisons test for **c**.

To examine a long-lasting protective effect of DA-HA, 10 µM DA-HA or 10 µM DA were treated every day from the beginning of rat primary neuronal culture for 14 days, and NeuN and MAP2 were monitored on day 14 (Fig. 4a-c). Daily counting of survived neurons for 14 days demonstrated that DA-HA treated group showed no significant neuronal death throughout the entire 14 days (Fig. 4d). In contrast, DA treated group showed a gradual and significant decrease in the percentage of survived neurons (Fig. 4d). On day 14, DA significantly reduced the percentage of survived neurons and NeuN^+^ neurons compared to control whereas DA-HA did not (Fig. 4e, f). These results indicated that, unlike DA, DA-HA has a long-lasting protective effect.

**Fig. 4.**
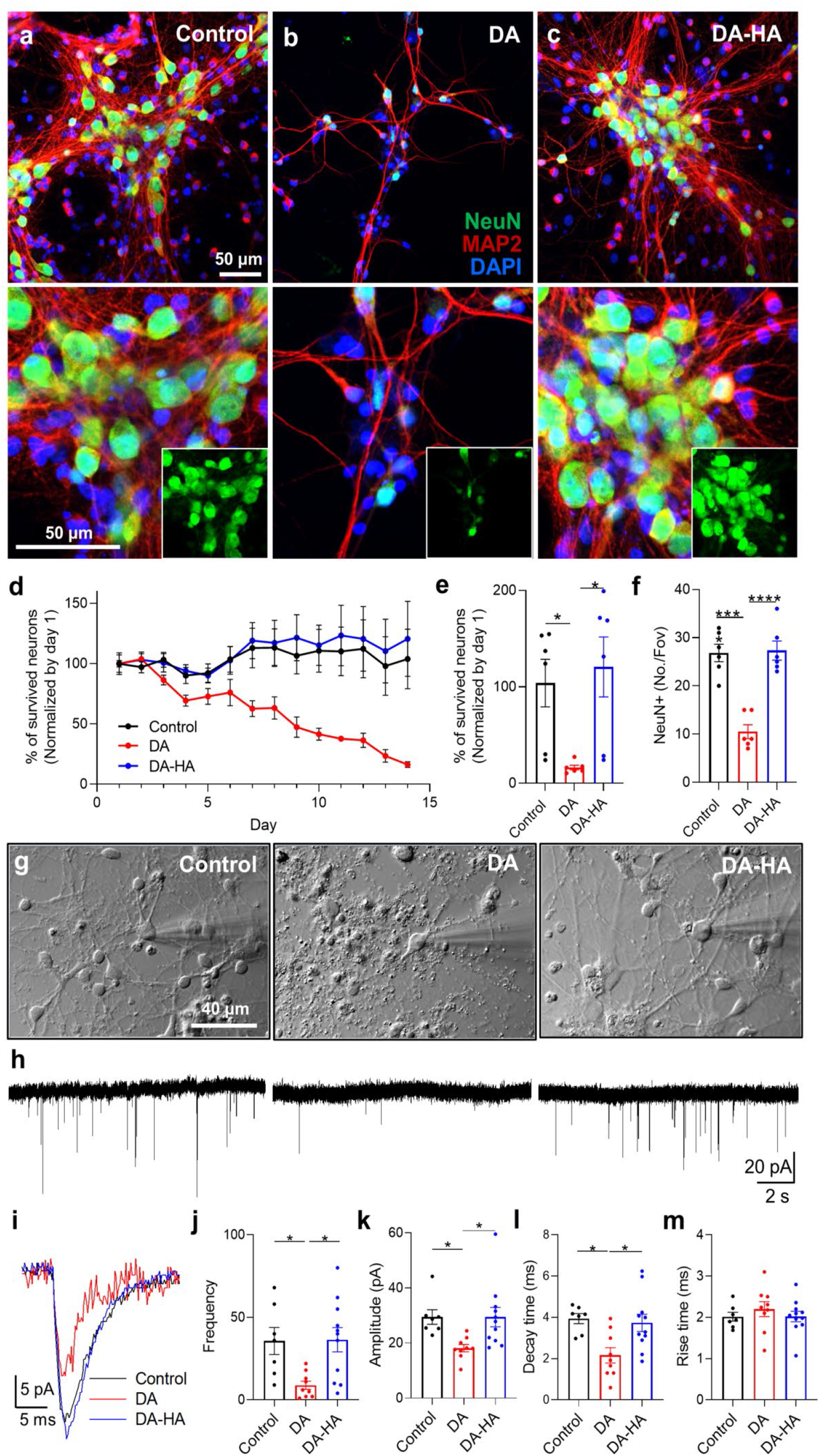
Long-term stability and safety of DA-HA on cultured neurons. **a-c** Representative images of NeuN (green), MAP2 (red), and DAPI (blue) of cultured neurons with no treatment (**a**), 10 µM DA (**b**), and DA-HA (**c**) for 14 days. Scale bar: 50 µm. Insets of lower pictures indicate NeuN^+^ cells. **d** Time-dependent changes of survived neurons under treatment of DA and DA-HA for 14 days (n = 6 for each condition). Two-way ANOVA, interaction between day & group F(26, 195) = 3.45, p < 0.0001; day F(13, 195) = 1.52, p = 0.11; group F(2, 15) = 7.05, p = 0.0069. **e** Summary bar graph of the percentage of survived neurons at day 14 from **d** (n = 6 for each condition). One-way ANOVA, F(2, 15) = 5.95, p = 0.01; Control vs DA, *p < 0.05; DA-HA vs DA, *p < 0.05. **f** Summary bar graph of the number of NeuN (n = 6 for each condition). One-way ANOVA, F(2, 15) = 29.10, p < 0.0001; Control vs DA, ***p < 0.001; DA vs DA-HA, ****p < 0.0001. **g, h** Bright-field images (**g**) and representative sEPSC traces (**h**) of cultured neurons under the treatment of vehicle (control, left), 10 µM DA (middle), and DA-HA (right) for day 14. **i** Superimposed traces of averaged sEPSC with no treatment (black), DA (red), and DA-HA (blue). **j** Summary bar graph of sEPSC frequency (n = 7, 9, and 11 for **j-m**). One-way ANOVA, F(2, 26) = 8.002, p = 0.002; Control vs DA and DA vs DA-HA, *p < 0.05. **k** Summary bar graph of sEPSC amplitude. One-way ANOVA, F(2, 24) = 5.13, p = 0.014; Control vs DA and DA vs DA-HA, *p < 0.05. **l** Summary bar graph of the rise time of sEPSC. One-way ANOVA, F(2, 24) = 0.51, p = 0.61. **m** Summary bar graph of the decay time of sEPSC. One-way ANOVA, F(2, 24) = 6.30, p = 0.0063; Control vs DA and DA vs DA-HA, *p < 0.05. Tukey’s multiple comparison test for **e, f, j, k**, and **m**.

To evaluate the long-lasting protective effect of DA-HA on synaptic transmission, we recorded spontaneous excitatory postsynaptic current (sEPSC) to measure glutamatergic synaptic activities (Fig. 4g, h). DA treated group showed significantly decreased sEPSC frequency and amplitude (Fig. 3j, k). In contrast, DA-HA treated group did not show a significant reduction in sEPSC frequency and amplitude (Fig. 4j, k). To scrutinize the release kinetics of glutamate, we measured the rise and decay time of each averaged sEPSC (Fig. 4i). We found that DA-HA did not show altered kinetics of sEPSC, whereas DA showed a significant decrease in decay time (Fig. 4l, m). These results indicated that DA-HA showed a long-lasting protective effect on synaptic transmission. Taken together, unlike DA, DA-HA shows long-term stability and safety on cultured neurons.

### Minimal extracellular autoxidation of DA-HA confers safety and stability

To investigate the molecular mechanism of DA-HA’s safety and stability, we considered two possible mechanisms involving intracellular ROS or extracellular oxidation. By using nomifensine, a DA transporter inhibitor, we eliminated the possible contribution of intracellular ROS in DA-induced cytotoxicity (Supplementary Fig. 2). To examine the contribution of extracellular oxidation, we established a simple assay system for the detection of ascorbic acid-sensitive autoxidation in culture media (Supplementary Fig. 3)^28^. Before investigating the relationship between autoxidation and cell viability, we assessed and compared the level of autoxidation (darkening of media color due to the formation of polydopamine, Fig. 1b) between DA and DA-HA in the neurobasal culture media only without neurons (Fig. 5a). We observed that both DA and DA-HA showed no significant autoxidation up to 10 µM concentration (Fig. 5b, c). At higher concentrations, DA-HA showed significantly less autoxidation than DA at both 24 and 72-h incubation times (Fig. 5b, c), implicating that DA-HA is highly resistant to autoxidation compared to DA.

**Fig. 5.**
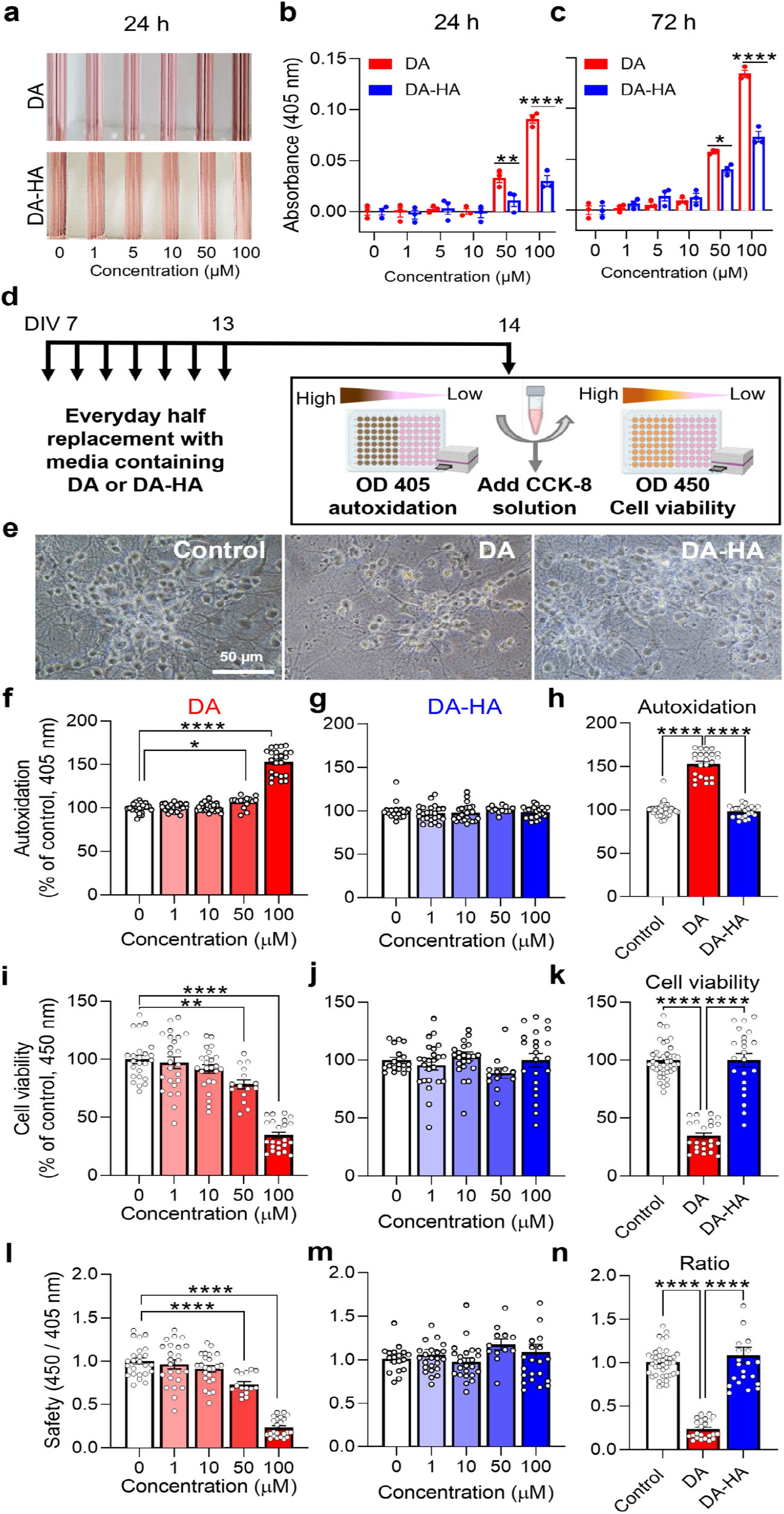
DA-HA shows virtually no toxicity, which is highly correlated with a low level of autoxidation. **a** Representative photographs for color change of neurobasal medium in the presence of 0, 1, 5, 10, 50, and 100 µM of DA and DA-HA for 24 hrs. **b** Summary bar graph of autoxidation in the presence of DA and DA-HA for 24 hrs (n = 3 for each condition). Two-way ANOVA, interaction between concentration & group F(5, 24) = 15.65, p < 0.0001; concentration F(5, 24) = 63.98, p < 0.0001; group F(1, 24) = 32.68, p < 0.0001; 50 µM DA vs DA-HA, **p < 0.01; 100 µM DA vs DA-HA ****P < 0.0001. **c** Summary bar graph of autoxidation in the presence of DA and DA-HA for 72 hrs (n = 3 for each condition). Two-way ANOVA, interaction between concentration & group F(5, 24) = 29.08, p < 0.0001; concentration F(5, 24) = 261.5, p < 0.0001; group F(1, 24) = 27.1, p < 0.0001; 50 µM DA vs DA-HA, *p < 0.05; 100 µM DA vs DA-HA ****P < 0.0001. **d** An experimental schedule for measuring autoxidation and cell viability of DIV 14 cortical cultured neurons with everyday supplementation of 1, 10, 50, and 100 µM DA and DA-HA for 7 days. **e** Bright-field images of control, DA, and, DA-HA treated neurons. Scale bar: 50 µm. **f** Summary bar graph of oxidation level in the presence of DA (n = 24, 24, 24, 15, and 24). One-way ANOVA, F(4, 106) = 181.5, p < 0.0001; 0 vs 50 µM, *p < 0.05; 0 vs 100 µM, ****p < 0.0001. **g** Summary bar graph of oxidation level in the presence of DA-HA (n = 18, 24, 23, 12, and 22). One-way ANOVA, F(4, 94) = 0.7763, p = 0.54. **h** Summary bar graph of oxidation level comparing 100 µM DA and DA-HA (n = 42, 24, and 22). One-way ANOVA, F(2, 85) = 237.2, p < 0.0001; Control vs DA and DA vs DA-HA, ****p < 0.0001. **i** Summary bar graph of cell viability in the presence of DA (n = 24, 24, 24, 15, and 23). One-way ANOVA, F(4, 105) = 50.88, p < 0.0001; 0 vs 50 µM, **p < 0.01; 0 vs 100 µM, ****p < 0.0001. **j** Summary bar graph of cell viability in the presence of DA-HA (n = 18, 24, 23, 12, and 22). One-way ANOVA, F(4, 94) = 1.23, p = 0.3024. **k** Summary bar graph of cell viability comparing 100 µM DA and DA-HA (n = 42, 23, and 22). One-way ANOVA, F(2, 84) = 109.1, p < 0.0001; Control vs DA and DA vs DA-HA, ****p < 0.0001. **l** Summary bar graph of the ratio of cell viability/autoxidation in the presence of DA (n = 24, 24, 24, 15, and 23). One-way ANOVA, F(4, 105) = 74.00, p < 0.0001; 0 vs 50 µM, ****p < 0.05; 0 vs 100 µM, ****p < 0.0001. **m** Summary bar graph of the ratio of cell viability/autoxidation in the presence of DA-HA (n = 18, 24, 23, 12, and 22). One-way ANOVA, F(4, 94) = 1.18, p = 0.3245. **n** Summary bar graph of safety (ratio of cell viability/autoxidation) comparing 100 µM DA and DA-HA (n = 42, 23, and 22). One-way ANOVA, F(2, 84) = 87.59, p < 0.0001; Control vs DA and DA vs DA-HA, ****p < 0.0001. Dunnett’s multiple comparisons test for **f, h, i, k, l**, and **n**.

To investigate the relationship between autoxidation and cell viability, we sequentially monitored autoxidation and cell viability from neuronal culture by measuring OD at 405 nm and 450 nm, respectively in the presence of DA-HA or DA for 7 days (Fig. 5d, e). To maintain a similar concentration of DA and DA-HA despite an active uptake through DA transporters, we replaced the culture media with half amount of fresh media daily for 7 days (Fig. 5d). Consistent with the media only measurements, we found that DA showed increased autoxidation in a concentration-dependent manner, whereas DA-HA did not (Fig. 5f, g). At 100 µM concentration, DA showed a significantly higher autoxidation than DA-HA, whereas DA-HA showed virtually no autoxidation compared to control (Fig. 5h). Consistent with autoxidation results, DA reduced cell viability in a concentration-dependent manner, whereas DA-HA did not (Fig. 5i, j). At 100 µM concentration, DA showed a significant reduction in cell viability, whereas DA-HA did not (Fig. 5k). To evaluate the degree of safety, we calculated the ratio of OD at 450 nm divided by OD at 405 nm for each well in the culture plate (Fig. 5l-n). DA-HA showed a high degree of safety compared to DA (Fig. 5l-n). Taken together, unlike DA, DA-HA showed virtually no cytotoxicity which is highly correlated with the absence of autoxidation, indicating a high degree of safety.

Based on the fact that autoxidation of DA is negatively correlated with cell viability, we hypothesized that the oxidized form of DA directly causes neuronal death. To test this hypothesis, we utilized DA and its derivatives with various degrees of autoxidation including L-DOPA and 6-OHDA (Fig. 1c, d). To determine the degree and time course of autoxidation, we assessed and compared the level of autoxidation in DA, L-DOPA, 6-OHDA, or DA-HA containing culture media (Fig. 6a). At 0 h, only 6-OHDA showed significantly higher baseline autoxidation (Fig. 6b, c). By the time of 48 h, all DA and its derivatives reached a maximal level of autoxidation and saturation at 72 h, which were all significantly higher than control (Fig. 6b, d). Unexpectedly, L-DOPA showed the highest autoxidation level, whereas DA-HA showed the lowest (Fig. 6d). The same pattern persisted even after 49-day incubation (Fig. 6e). Based on these results, we defined “pre-autoxidized form” or “pre-autoxidation” as DA or its derivatives in its autoxidized state after 48-h incubation.

**Fig. 6.**
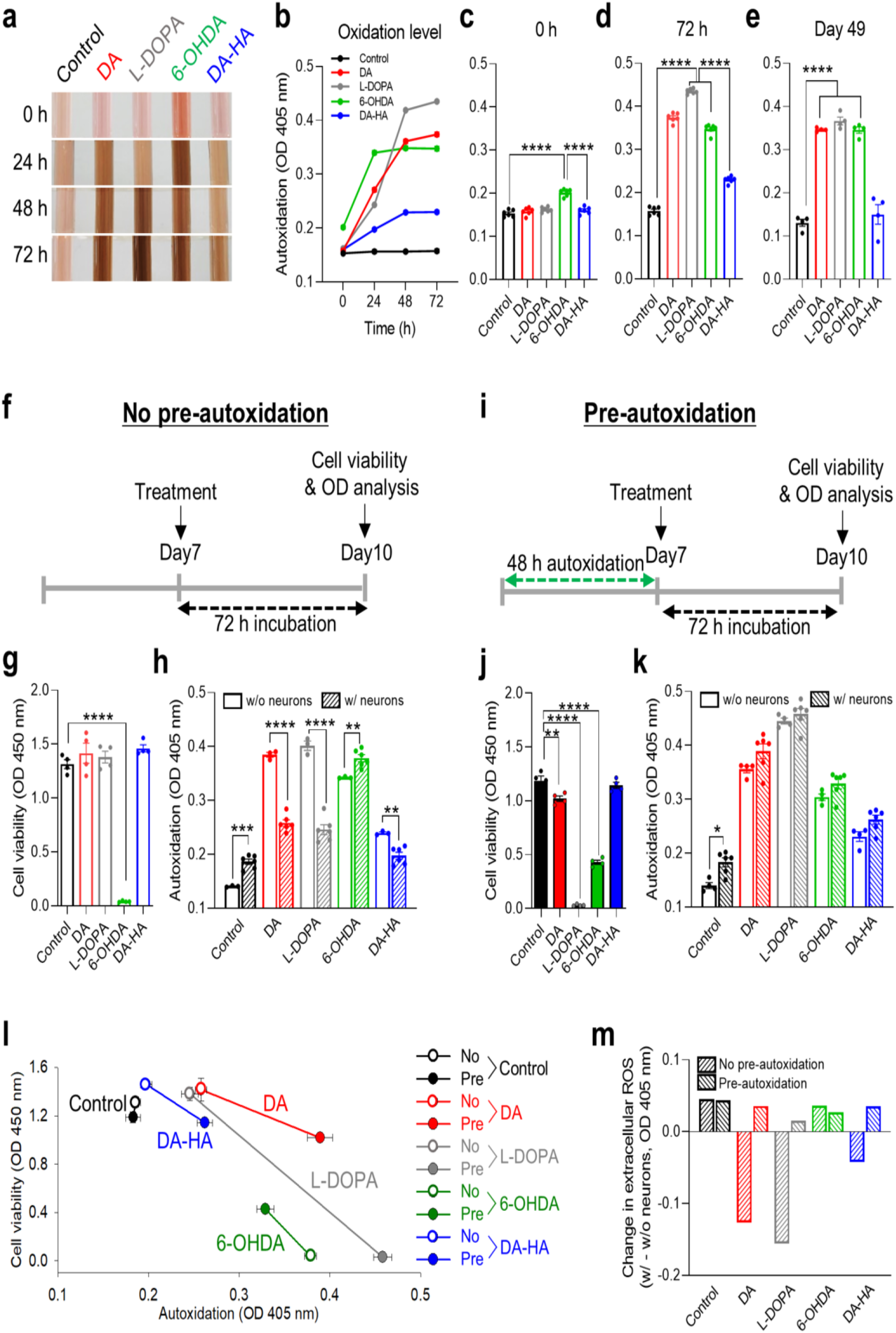
Minimal extracellular autoxidation is the key molecular factor for safety and stability. **a** Representative photographs of the color change of neurobasal medium treated with 200 µM DA, L-DOPA, 6-OHDA, and DA-HA at 0, 24, 48, and 72 h. **b** Summary for autoxidation level of 200 µM DA, L-DOPA, 6-OHDA, and DA-HA at 0, 24, 48, and 72 h (n = 6 for each condition). **c** Summary bar graph of autoxidation at 0 h (n = 6 for each condition). One-way ANOVA, F(4, 25) = 38.04, p < 0.0001; Control vs 6-OHDA, ****p < 0.0001. **d** Summary bar graph of autoxidation at 72 h (n = 6 for each condition). One-way ANOVA, F(4, 25) = 889.9, p < 0.0001; Control vs DA and DA-HA vs DA, ****p < 0.0001. **e** Summary bar graph of autoxidation at day 49 (n = 4 for each condition). One-way ANOVA, F(4, 15) = 96.13, p < 0.0001; Control vs DA, Control vs L-DOPA, and Control vs 6-OHDA, ****p < 0.0001. **f** Experimental schedule for cell viability and oxidation level with no autoxidation. **g** Summary bar graph of cell viability with no autoxidation (n = 4 for each condition). One-way ANOVA, F(4, 15) = 127.7, p < 0.0001; Control vs 6-OHDA, ****p < 0.0001. **h** Summary bar graph of oxidation level with and without neuronal cells with no autoxidation condition (n = 6 and 3). Unpaired t-test, Control w/ vs w/o, ***p < 0.001; DA w/ vs w/o and L-DOPA w/ vs w/o, ****p < 0.0001; 6-OHDA w/ vs w/o and DA-HA w/ vs w/o, **p < 0.01. **i** Experimental schedule for cell viability and oxidation level with 48-h autoxidation. **j** Summary bar graph of cell viability with autoxidation of 200 µM DA, L-DOPA, 6-OHDA, and DA-HA for 48 h (n = 4 for each condition). One-way ANOVA, F(4, 15) = 338.2, p < 0.0001; Control vs DA, **p < 0.01; Control vs L-DOPA and Control vs 6-OHDA, ****p < 0.0001). **k** Summary bar graph of autoxidation level with and without neuronal cells with 48-h autoxidation condition (n = 6 and 4). Unpaired t-test, Control w/ vs w/o cell, *p < 0.05. **l** The relationship between cell viability and autoxidation from **f**-**k. m** Summary bar graph of uptake level in no pre-autoxidation and pre-autoxidation conditions (n=1). Dunnett’s multiple comparisons test for **c, e, g**, and **j**. Tukey’s multiple comparison test for **d**.

Next, to directly examine whether pre-autoxidation is critical for cytotoxicity, we sequentially measured cell viability and autoxidation level in the absence or presence of the pre-autoxidized form of DA and its derivatives (Fig. 6f, i). At the same time, to assess the possible contribution of neuronal uptake to cytotoxicity, we compared autoxidation with or without neurons. In no pre-autoxidation condition, only 6-OHDA showed an almost complete reduction of cell viability (Fig. 6g), consistent with the high baseline autoxidation. In pre-autoxidation condition, DA and L-DOPA showed a significant decrease in cell viability, with an almost complete reduction with L-DOPA (Fig. 6j), raising a possibility that autoxidation of DA and L-DOPA directly causes neuronal death. Intriguingly, 6-OHDA showed a significant improvement in cell viability under pre-autoxidation condition (Fig. 6l), suggesting that, unlike DA and L-DOPA, 6-OHDA itself is toxic but the oxidized form of 6-OHDA might be less toxic. In contrast, DA-HA showed no cytotoxicity regardless of pre-autoxidation (Fig. 6g, j, l). These results indicate that at least with DA and L-DOPA, pre-autoxidation is critical for cytotoxicity.

We then compared autoxidation (extracellular ROS level) with or without neurons to indirectly assess the possible contribution of neuronal uptake to cytotoxicity (Fig. 6h, k). Under no pre-autoxidation condition, the autoxidation level of the control group with neurons was significantly higher than without neurons (Fig. 6h), suggesting that cell metabolism causes higher oxidation ^29^. In contrast, DA and L-DOPA showed significantly lower autoxidation with neurons than without neurons (Fig. 5h, m), suggesting that DA and L-DOPA are actively taken up by neurons. In contrast, in the pre-autoxidation condition, DA and L-DOPA showed no significant autoxidation level between with and without neurons (Fig. 6k, m), suggesting that there was a minimal uptake of the pre-autoxidized form of DA or L-DOPA by neurons. DA-HA showed slightly but significantly higher autoxidation in with neurons under the no pre-autoxidation condition but not under the pre-autoxidation condition (Fig. 6h, k, m). This could be due to released DA from DA-HA ^22^ or the presence of a trace amount of free DA in DA-HA. These results indicate that uptake of DA and L-DOPA does not contribute to cytotoxicity. Instead, it alleviates cytotoxicity by removing extracellular DA and L-DOPA. Taken together, the minimal autoxidation level of DA-HA is the key molecular factor for its safety and stability.

### DA-HA does not induce dopaminergic neuronal death in substantia nigra

To investigate whether DA-HA retains its safety and stability *in vivo*, we directly injected DA-HA into mouse substantia nigra pars compacta (SNpc, Fig. 7a). After 14 days of injection of saline, 5 mM DA, 5 mM HA, 5 mM DA-HA, 5mM 6-OHDA or 50 mM 6-OHDA, mice were sacrificed and immunostaining of tyrosine hydroxylase (TH, Fig. 7b) was performed to measure dopaminergic neurons in SNpc (cell body) and striatum (axonal projection). We found that the TH immunofluorescence ratio (ipsilateral divided by contralateral side) showed no significant difference between DA-HA- and saline-injected groups in both SNpc and striatum regions (Fig. 7b, c), indicating that DA-HA shows no toxicity to dopaminergic neuron in SNpc area. Similar results were obtained with DA and HA (Fig. 7b and c). In contrast, 6-OHDA showed a significant decrease in TH immunofluorescence in a dose-dependent manner (Fig. 7c). Taken together, unlike 6-OHDA, DA-HA does not cause dopaminergic neuronal death *in vivo*, substantiating its *in vivo* safety.

**Fig. 7.**
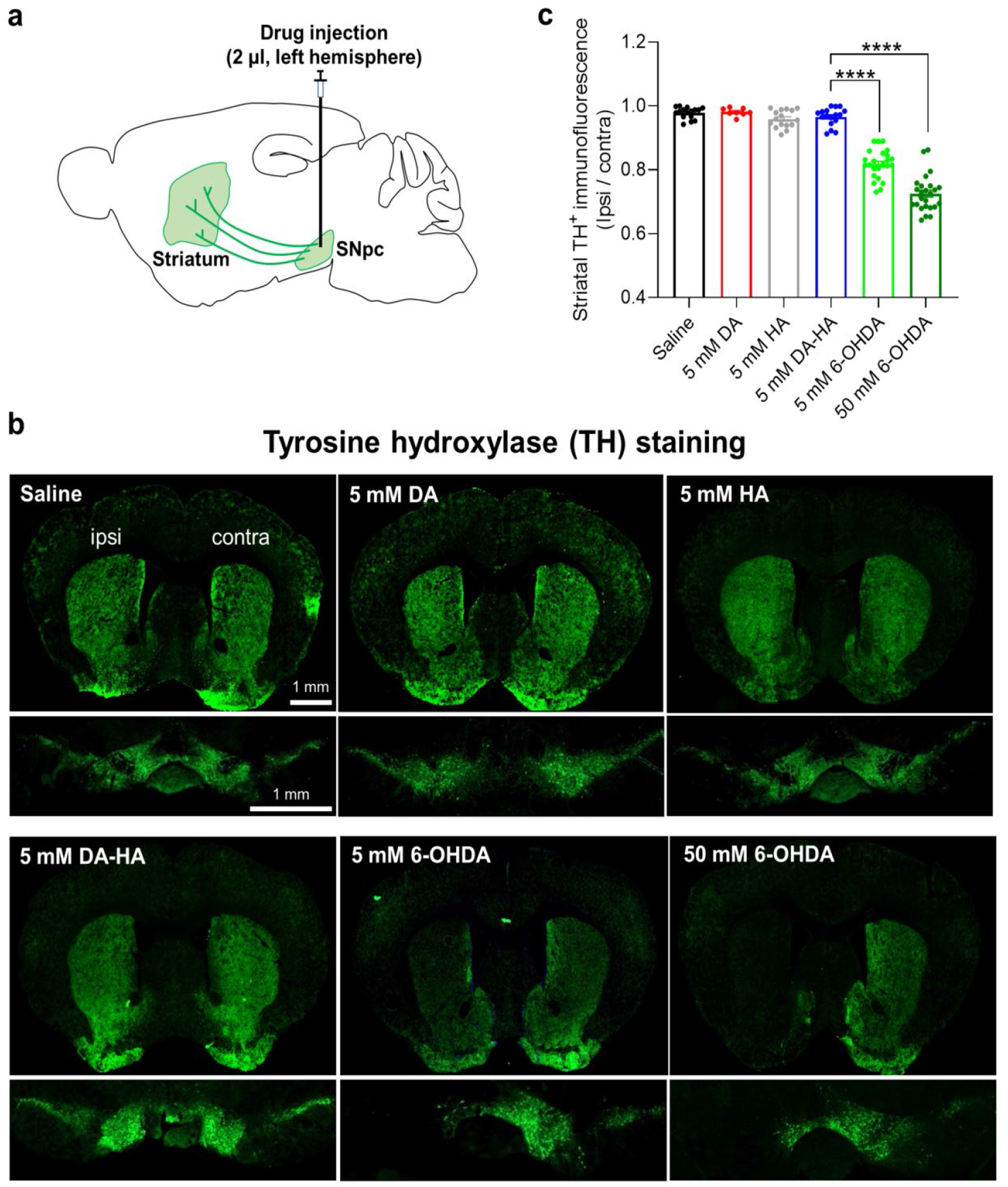
DA-HA does not induce dopaminergic neuronal death *in vivo*. **a** A schematic diagram of drug injection into SNpc. **b** Representative fluorescent images of TH staining in the striatum (top) and SNpc (bottom) with the injection of saline, 5 mM DA, HA, DA-HA, 6-OHDA, and 50 mM 6-OHDA into SNpc. Scale bar: 1 mm. **c** Summary bar graph of optical density of TH^+^ fluorescence (n = 16, 8, 16, 16, 24, and 24 of slices). One-way ANOVA, F(5, 100) = 4.699, p < 0.0001; 5 mM DA-HA vs 5 mM 6-OHDA and 5 mM DA-HA vs 50 mM 6-OHDA, ****p < 0.0001. Tukey’s multiple comparison test for **c**.

### DA-HA alleviates apomorphine-induced rotational behavior in 6-OHDA Parkinson’s disease model

Finally, to test whether DA-HA can exert a beneficial effect on parkinsonian motor symptoms, we employed the 6-OHDA-induced mouse model of Parkinson’s disease ^30^. We injected 6-OHDA (2 µl, 15 µg, left hemisphere only) into SNpc, followed by bilateral insertions of guide cannula into the striatum (Fig. 8a, b). We performed an apomorphine-induced rotation test 14 days after surgery (Fig. 8a). We found that the striatal infusion of 2 µl for 10 min of either 2.5 mM DA-HA or 2.5 mM DA into the left hemisphere caused a significant reduction of apomorphine-induced counterclockwise rotation within 10 min after drug infusion (Fig. 8c and d), suggesting that DA-HA significantly restored the dopamine signaling in the striatum of the ipsilateral side of 6-OHDA lesion. In contrast, the same dosage of L-DOPA showed no significant change in apomorphine-induced counterclockwise rotation within 10 min after drug infusion (Fig. 8c, d), indicating that the beneficial effect of L-DOPA is not immediate. Taken together, these results strongly suggest that DA-HA can immediately alleviate parkinsonian motor symptoms in an animal model of Parkinson’s disease.

**Fig. 8.**
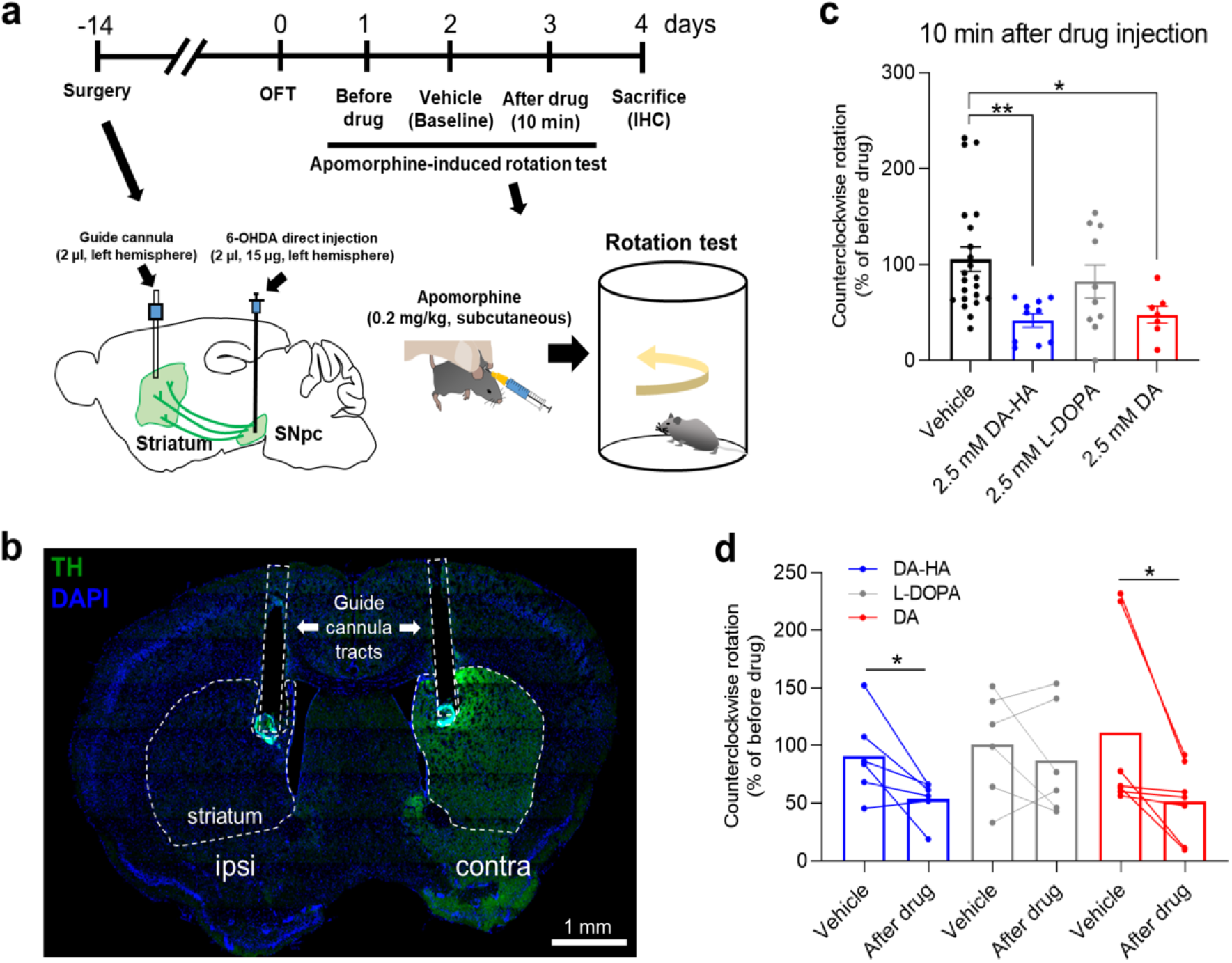
DA-HA alleviates apomorphine-induced rotational behavior in 6-OHDA Parkinson’s disease model. **a** Experimental schedule and detailed schematic diagrams for surgery and apomorphine-induced rotation test. Apomorphine was delivered by subcutaneous injection (s. c.) **b** Representative TH^+^ fluorescent image of striatum for validation of guide cannula location. Dotted lines indicate guide cannula tracts and striatum. Scale bar: 1 mm. **c** Unpaired comparison of apomorphine-induced rotation test with 10 m after injection of vehicle (0.02% ascorbic acid), 2.5 mM DA-HA, L-DOPA, and DA (n = 22, 10, 10, 7) into the striatum. One-way ANOVA, F(3, 45) = 5.05, p < 0.0042; Vehicle vs DA-HA, **p < 0.01; Vehicle vs DA, *p < 0.05. **d** Paired comparison between vehicle, 10 m, and 24 h after drug injection (n = 6, 6, 7). Paired t-test, Vehicle vs DA-HA and Vehicle vs DA, *p < 0.05. Tukey’s multiple comparison test for **c**.

## Discussions

In this study, we have demonstrated for the first time that DA-HA can be utilized for brain-related applications. This is possible because DA-HA possesses similar pharmacological and functional properties as DA. More importantly, unlike DA and L-DOPA, DA-HA shows virtually no autoxidation, which is highly correlated with low neurotoxicity. Mechanistically, we have demonstrated that DA-HA’s safety and stability are attributed to its minimal extracellular autoxidation. We have further recapitulated DA-HA’s functionality and safety *in vivo* as we discovered that DA-HA does not induce dopaminergic neuronal death in mouse SNpc and that DA-HA alleviates parkinsonian motor symptoms in the 6-OHDA mouse model of Parkinson’s disease. Based on these findings, we propose that DA-HA with DA-like functionalities and minimal toxicity can be an effective therapeutic substitute for L-DOPA in Parkinson’s disease.

How does DA-HA manifest similar functional properties as DA? Here we found that DA-HA can directly activate DA receptors and evoke a Ca^2+^ response in NAc astrocytes (Fig. 2j-l). How can DA-HA activate endogenous DA receptors? Previous structural studies reported that both hydroxyl (-OH) and amine (-NH2) groups are critical for DA to interact with D^1^R and D^2^R ^31,32^. However, unlike DA, DA-HA has only free hydroxyl groups, but not a free amine group, which is conjugated to the carboxyl group of HA (Fig. 1a). Our study demonstrates that despite the lack of a free amine group, DA-HA can still activate DA receptors, implicating that the amine group is not critical for DA-HA to bind to DA receptors (Fig. 2). In support of this idea, it is known that other DA receptor agonists, such as SKF-82958 and SKF-83959, activate D1R (dopamine receptor type 1) in the absence of free amine group <^33,34^. In addition, DA-HA is composed of a long polymer chain, which might interfere with a proper interaction between DA-HA and DA receptor due to a possible steric hindrance (Fig. 1a). However, our results demonstrate that the long polymer chain of DA-HA does not appear to interfere with a proper interaction between DA-HA and DA receptors for activation of DA receptors. Despite the fact that DA-HA has critical structural differences compared to DA, DA-HA can activate an exogenous GRAB^DA2m^ sensor, which was originally developed by using D^2^R motif ^35^(Fig. 2g-i). Moreover, DA-HA evokes Ca^2+^ transients in NAc astrocytes through the activation of endogenous D^1^R in astrocytes ^27^(Fig. 2j-l), implying that DA-HA is able to bind to both D^1^R and D^2^R. Taken together, our study is the first report that DA-HA can act as an agonist of DA receptors, paving a new way for various brain-related applications.

Our study provides compelling lines of evidence that DA-induced cytotoxicity is majorly caused by extracellular autoxidation, but not by intracellular ROS through DA uptake. If intracellular ROS production (following DA uptake) is the major cause of DA-induced cytotoxicity, then inhibition of DA transporter should decrease DA-induced neuronal cell death. Contrary to this expectation, we found that inhibition of DA transporter increased (rather than decreased) the DA-induced neuronal cell death (Supplementary Fig. 2). Furthermore, we found that uptake of DA or L-DOPA by neurons does not contribute to cytotoxicity (Fig. 6g, j, l), but instead, it alleviates cytotoxicity by removing extracellular DA and L-DOPA (Fig. 6h, m). These results strongly suggest that extracellular autoxidation is the major factor for DA-induced cytotoxicity. In terms of DA-HA, we expected that DA-HA would not be taken up by neurons due to its long and bulky polymeric structure (Fig. 1a). Contrary to our expectation, DA-HA showed a slight but significant contribution of uptake in alleviating cytotoxicity (Fig. 6l, m). This could be due to an uptake of slowly released free DA or free DA-HA monomer from DA-HA polymer ^22^, or the presence of a trace amount of free DA in DA-HA. Nevertheless, this contribution by DA-HA uptake is a minor and negligible portion compared to DA and L-DOPA (Fig. 6m). Interestingly, contrary to DA, L-DOPA, and DA-HA, 6-OHDA resulted in reduced cytotoxicity after pre-autoxidation compared no pre-autoxidation condition (Fig. 6j). This opposite result suggests that 6-OHDA displays potent cytotoxicity regardless of the degree of autoxidation and that the 6-OHDA-induced cytotoxicity requires uptake of the un-oxidized form of 6-OHDA. This raises a unique possibility that the 6-OHDA-induced cytotoxicity is majorly caused by intracellular ROS after uptake, which is consistent with previous reports demonstrating that 6-OHDA-induced cytotoxicity was blocked by nomifensine treatment ^36,37^. Taken together, while DA, L-DOPA, and 6-OHDA exert severe cytotoxicity due to autoxidation or uptake, DA-HA exhibits minimal cytotoxicity due to the lack of autoxidation, which confers its long-term stability and safety.

What could be the extracellular molecular factor for DA- and L-DOPA-induced cytotoxicity? During the autoxidation process of DA, highly reactive DA metabolites such as dopamine-o-quinone, aminochrome, and 5,6-dihydroxyindole are produced (Fig. 1b). In addition, each oxidation step liberates two protons and two electrons as by-products (Fig. 1b). L-DOPA goes through a similar process (Fig. 1c). Liberated two protons should decrease extracellular pH, whereas liberated two electrons generate free radicals and superoxide. The consequence of prolonged extracellular acidosis is that it can cause neuronal and glial death ^38^, possibly via acidosis-mediated excessive efflux of chloride ions to neutralize extracellular pH. Consistent with this idea, we indeed observed cell shrinkage during prolonged DA treatment (Fig. 5e). Another possible mechanism of DA-induced cytotoxicity is the non-specific formation of toxic cysteinyl catechols between the membrane proteins and highly reactive DA metabolites, which are produced during autoxidation process ^39,40^. Unlike DA and L-DOPA, DA-HA lacks the autoxidation process which results in minimal cytotoxicity. Consistently, prolonged DA-HA treatment did not induce cell shrinkage (Fig. 5e). Taken together, we have identified several molecular factors for DA- and L-DOPA-induced cytotoxicity, including the formation of cysteinyl catechols, extracellular acidosis, and ROS generation.

Based on our findings, we propose that DA-HA can be an effective therapeutic substitute to replace L-DOPA. The current understanding of how L-DOPA alleviates parkinsonian motor symptoms is through upregulation of DA synthesis and replenishment of depleted DA to activate DA receptors ^41,42^. For 70 years since the first clinical report, L-DOPA has been dominating the clinical field of Parkinson’s disease as a “gold standard” drug, even though it cannot completely cure the disease. Instead, L-DOPA effectively relieves the parkinsonian motor symptoms for a short period of time, while patients take L-DOPA 3 or 4 times daily ^43^. Unfortunately, L-DOPA has multiple adverse side-effects such as nausea, dizziness, headache, and somnolence ^43^. On top of these acute side-effects, chronic L-DOPA administration causes a severe side-effect called levodopa-induced dyskinesia ^44^. Therefore, there has been a long-standing and desperate need to develop a new approach to overcome the drawbacks of this old medication. DA-HA shows minimal cytotoxicity compared to L-DOPA while maintaining DA-like functionalities, ensuring minimal acute side-effects. Unlike L-DOPA, DA-HA is most likely resistant to uptake due to its bulky polymeric structure. This can be an important advantage of DA-HA over L-DOPA because chronic use of L-DOPA is known to increase intracellular oxidative stress through monoamine oxidase-led enzymatic degradation of synthesized DA, causing neuronal damage and cytotoxicity ^45^. Another key advantage of using DA-HA is that it activates DA receptors directly from outside of the cell, whereas L-DOPA requires multiple steps: uptake, enzymatic conversion to DA, release into extracellular space, and DA receptor activation. This advantage of DA-HA over L-DOPA ensures minimal chance for confounding effects. A previous study reported that free DA from DA-HA can be slowly released by endogenous extracellular hyaluronidase ^46^. This can ensure the long-lasting effect of DA-HA with just a single administration. One potential drawback of DA-HA is that, unlike L-DOPA, DA-HA is most likely BBB-impermeable ^47^. Future investigations are needed to develop DA-HA derivatives with high BBB-permeability.

In conclusion, our study overcomes the limitations of currently available DA-mimetics by unveiling the disguised characteristics of DA-HA: 1) DA-HA shows long-lasting stability and safety with DA-like functionalities, 2) DA-HA is most likely resistant to uptake, 3) DA-HA activates DA receptor directly from outside of the cell, and 4) DA-HA can be slow acting. The new concepts and tools we have developed in this study will prove useful in developing better DA-mimetics to treat DA-depleted neurological disorders such as Parkinson’s disease.

## Methods

### Synthesis of DA-HA

DA-HA was synthesized as a previous report ^48,20^. 1 g of HA was dissolved in 100 mL of phosphate-buffered saline (PBS) solution and the pH was adjusted to 5.5 with a hydrochloric acid (HCl) aqueous solution. The solution was purged with nitrogen for 30 minutes. Then, 338 mg of EDC and 474 mg of DA were added and the pH of the reaction solution was maintained at 5.5 for 2 h. Unreacted chemicals and urea by-products were removed by extensive dialysis, and afterward, the conjugate was lyophilized. In order to avoid oxidation, the conjugate was stored at 4 °C in a desiccator under a vacuum and protected from light. In this study, DA-HA concentration was considered as DA concentration of DA-HA. In this study, we used 0.5 and 1 of the feed molar ratio of DA-HA (Table 1).

### Ultraviolet (UV) Spectrophotometry and Nuclear Magnetic Resonance (NMR) Analysis of DA-HA

The degree of substitution of DA in the conjugate was determined using a UV-vis spectrophotometer (Shimadzu UV-2501 PC) and 1 cm quartz cells. A solution of 1 mg / mL in 0.15 M sodium chloride (NaCl) was prepared for the UV analysis. The samples were dissolved in deuterated water for 24 h at concentrations of 1 mg/mL and 10 mg/mL for ^1^H-NMR and ^13^C-NMR analyses, respectively. The results were obtained using a spectrometer (Varian Unity plus 300) and tetramethylsilane (TMS) as the internal standard. The spectra were recorded at 298 K and 300 MHz for 1 H-NMR analysis and 333 K and 75.4 MHz for ^13^C-NMR analysis.

### Subject

Postnatal day (P)1 of Sprague Dawley (SD) rats for the primary cultured neurons were purchased (OrientBio, Korea). Male and female adult C57BL/6J mice were kept on a 12 h light-dark cycle with controlled temperature and humidity and had ad libitum access to food and water (IBS Research Solution Center). Mice were housed in a common cage (3 - 5 mice/cage) and housed in individual housing after guide cannula implantation surgery. All experiments were conducted in accordance with animal study protocols approved by the Institutional Animal Care and Use Committee of the Korea Institute of Toxicology (KIT, Daejeon, Korea) and the Institute for Basic Science (IBS, Daejeon, Korea). Animal care was performed following National Institutes of Health (NIH) guidelines.

### Rat primary cortical astrocyte culture

Rat primary cortical astrocytes were prepared from P1 of SD rats as described ^49^. The cerebral cortex of the rat was dissected free of adherent meninges, minced, and dissociated into single-cell suspension by trituration. Astrocytes were grown in Dulbecco’s modified Eagle’s medium (DMEM, Gibco, USA) supplemented with 10% heat-inactivated horse serum, 10 % heat-inactivated fetal bovine serum, and 1% penicillin-streptomycin (100 units/ml penicillin–0.1 mg/ml streptomycin). Cultures were maintained at 37°C in a humidified 5% CO^2^ incubator. On the days *in vitro* (DIV) 3, cells were vigorously washed with repeated pipetting and the media was replaced to get rid of debris and other floating cell types. On DIV 7-8, cells were replated onto coverglass (8×10^4^ per well) coated with 50 µg/ml Poly-D-Lysine (PDL, Merck, USA) for imaging experiments.

### GRABDA2m transfection

On DIV 7-8, when astrocytes were trypsinized and collected for the transfection, 1.3×10^6^ primary cultured astrocytes were transfected with 4 µg AAV-GFAP104-GRAB^DA2m^ with an optimized electroporation method (1300V, 20 pulse width, 2 pulses) using the electroporator (Neon Transfection System, Korea). 8×10^4^ transfected cells were loaded onto a cover glass of each well and were incubated for 24 - 36 h.

### In vitro GRABDA2m imaging and analysis

External solution contained (mM): 150 NaCl, 10 HEPES, 3 KCl, 2 CaCl^2^, 2 MgCl^2^, 5.5 glucose; pH adjusted to pH 7.3 - 7.4 and osmolarity to 325 - 330 mOsm/kg. Intensity images of 510 nm wavelength were taken from excitation wavelength at 450 ± 20 nm using Electron Multiplying Charge-Coupled Device (EMCCD) camera (Ixon 867, Andor Oxford instrument). The astrocytes were incubated for 24 h to express GRAB^DA2m^ sensor. Once an astrocyte is targeted, DA and DA-HA were treated and peak amplitudes of GRAB^DA2m^ transients were measured. For measuring concentration-dependency, peak amplitudes of GRAB^DA2m^ transients were measured under treatment of 0.01, 0.1, 0.5, and 1 µM DA or DA-HA. Images are acquired for every 1 s.

### Virus construct

The AAV-hSyn-GRAB^DA2m^ viral vector was kindly provided by Yulong Li’s laboratory. This vector was subcloned into AAV-GFAP104-GRAB^DA2m^ for astrocytic expression. AAV containing GFAP104-GRAB^DA2m^ (titer = 5.32 × 10^12^) and GfaABC1D-GCaMP6f (titer = 4.09 × 10^12^) were packaged by the IBS virus facility (IBS, Daejeon, South Korea). Viruses were diluted immediately before injection (1/3 dilution with saline).

### Stereotaxic injection

Mice were anesthetized with isoflurane (3% for induction and 1.5% during surgery) and moved to a stereotaxic apparatus (68537, RWD Life Science). After shaving and cutting the skin of the mouse head, holes were made bilaterally in the skull to target the nucleus accumbens (NAc) using the following coordinates ^27^: AP, 1.5 mm; ML, ±0.75 mm; DV, −4.5 mm from the bregma. The virus was prepared and loaded into a vertically pulled VWR microdispenser (53508–375, SP Scientific, PA, USA) and was injected at a rate of 0.1 μL/ min with a total of 0.5 μL volume in each hemisphere using a micro-infusion pump (Legato 130, KD Scientific). For toxicity test *in vivo*, DA (H8502, Sigma Aldrich), HA, DA-HA, and 6-OHDA (2547, Tocris) were injected into the left hemisphere of substantia nigra pars compacta (SNpc) using the following coordinates: AP, -3.3 mm; ML, 1.3 mm; DV, -4.2 mm from the bregma. After termination of injection, let the injection needle stay for an additional 10 min before the withdrawal. The body temperature of the mouse was kept at 37 °C during injection (ThermoStar Homeothermic Monitoring System, RWD). After at least 2 weeks of recovery, the mice were used for imaging experiments. AAV-GFAP104-GRAB^DA2m^ and AAV-GfaABC1D-GCaMP6f were used for DA and Ca^2+^ imaging, respectively.

### Acute brain slice preparation

Mice were deeply anesthetized with 3 % isoflurane, followed by decapitation. The brain was quickly removed from the skull and submerged in ice-cold dissection buffer (in mM): 212.5 sucrose, 26 NaHCO^3^, 10 D-glucose, 5 MgCl^2^, 3 KCl, 1.25 NaH^2^PO^4^ and 0.1 CaCl^2^; pH 7.4, saturated with 95% O^2^ and 5% CO^2^. 300-μm-thick coronal slices were prepared by a vibratome (D.S.K LinearSlicer pro 7, Dosaka EM Co. Ltd, Kyoto, Japan). Slices were then transferred to an incubation chamber with artificial cerebrospinal fluid (aCSF) (in mM): 130 NaCl, 24 NaHCO^3^, 3.5 KCl, 1.25 NaH^2^PO^4^, 1.5 CaCl^2^, 1.5 MgCl^2^ and 10 glucose; pH 7.4, saturated with 95% O^2^ and 5% CO^2^, at room temperature. All imaging experiments were performed after 1 h of stabilization.

### Ex vivo DA imaging

Brain slices were transferred into a recording chamber where aCSF continuously flowed over slices at ∼1 mL/min. To visualize the GRAB^DA2m^ sensor, a blue LED (pE340fura, CoolLED, Hampshire, UK) was applied to brain slices under a fluorescent upright microscope (Zeiss Examiner.D1, NY, USA) with a 63x water objective (Zeiss, NY, USA). HA (10 µM), DA-HA (10 µM), and DA (10 µM) were treated by bath application controlled by a valve controller (VC-6 six channel valve controller, Warner instrument). To measure DA response from the GRAB^DA2m^ sensor, the peak of GRAB^DA2m^ transients during drug treatment was normalized by its baseline (ΔF/F^0^ ratio). ROIs were chosen based on the response from DA treatment. Image acquisition and ROI analysis were performed using Imaging Workbench (INDEC Biosystems, CA, USA) and ImageJ (NIH, MD, USA).

### Ex vivo calcium imaging

The procedure of calcium imaging was the same as DA imaging except for a little difference in virus and analysis. To monitor Ca^2+^ transients in NAc astrocytes, AAV-GfaABC1D-GCaMP6f virus was injected into NAc. Instead of measuring peak Ca^2+^ response (Due to the high amplitude of spontaneous Ca^2+^ response), we measured the area under the curve (AUC) during drug treatment.

### Rat primary cortical neuron culture

For cytotoxicity assay by counting of neuronal cells, primary cortical cultured neurons were prepared from mouse P0-P1 as described ^50^. Cortex was dissected and incubated with 0.25% trypsin for 10 min at 37°C. Trypsin was inactivated by adding DMEM containing 10% FBS. The media was decanted and Neurobasal media (Gibco, USA) consisted of 2 % B-27™ supplement (Gibco, USA), 1% L-glutamine (2mM, Gibco, USA), and 1% penicillin-streptomycin (100 units/ml penicillin–0.1 mg/ml streptomycin, Gibco, USA). The cell suspension was passed through a fire-polished Pasteur pipette rigorously and was repeated 3-5 times. Cells passed through a cell strainer with a pore size of 100 µm (SPL, Seoul, Korea) and 40 µm (SPL, Seoul, Korea) to remove non-triturated tissue. Cells were plated at 2.5× 10^5^ per each well of a 24-well plate and 8 × 10^4^ per each well of a 96-well plate. Cultures were maintained in a humidified atmosphere containing 5 % CO^2^ and 95 % O^2^ at 37 °C for 14 - 16 days.

### Immunocytochemistry

Cultured neurons on the coverslip were fixed overnight in 4 % paraformaldehyde (PFA). Neurons were washed with Phosphate-Buffered Saline (PBS) 3 times and incubated with a blocking solution for 1 h (0.5% Triton X-100, 1% BSA, 5% normal goat serum in PBS) at room temperature (RT). Primary antibodies in the blocking solution were incubated overnight at 4°C. Primary antibodies were diluted to the following amounts: anti-NeuN (Merck Millipore, MA, USA) 1:1000 and anti-MAP2 (Abcam, CB, UK) 1:1000. After washing with PBSTB (0.5% Triton X-100, 1% BSA in PBS) 2 times for 10 m, cells were incubated with secondary antibodies in PBSTB for 1 h at RT. Secondary antibodies were diluted to the following amounts: Alexa fluor 488 goat anti-guinea pig 1:1000, Alexa fluor 594 goat anti-rabbit 1:1000 (Invitrogen, CA, USA). And washing with PBST for 10 m 2 times and PBS for 10 m 1 time at RT, neurons on coverslips were incubated with a mounting medium including DAPI (VECTASHIELD, CA, USA) and mounted onto a slide glass. A series of fluorescent images were obtained by confocal microscope (Olympus, Tokyo, Japan) and analyzed by Image J software.

### Series cell counting assay

To assess the toxicity of DA-HA and DA, neurons were counted every day. Neurons were half-replaced with fresh media containing 20 µM DA and DA-HA every day. The number of neurons was counted every day for 14 days. The number of neurons was measured from 3 fields of view and averaged to obtain a single value. Day 1 indicated the starting day of surgical dissection.

### sEPSC measurement from rat primary cortical neurons

Whole-cell patch recording from cultured cortical neurons under voltage clamp (holding potential –70 mV) was acquired with a MultiClamp 700B amplifier digitized by Digidata 1322A data acquisition system (Molecular Devices, USA). The recording chamber was continually perfused with a recording solution composed of (mM): 150 NaCl, 3 KCl, 2 CaCl^2^, 5.5 glucose, and 10 HEPES (pH 7.3-7.4 by NaOH; osmolarity adjusted to 315–320 mOsm/kg with sucrose). Recording electrodes (4 – 7 MΩ) were filled with (mM): 150 CsMeSO^4^, 10 NaCl, 0.5 CaCl^2^, 10 HEPES, (pH adjusted to 7.3-7.4 with CsOH and osmolarity adjusted to 310-330 mOsm/kg with sucrose),0.5 mM QX 314. All electrophysiological data from cultured cells in this study were kept by a temperature controller (Warner instrument). Mini Analysis (Synaptosoft) was used to analyze the frequency, amplitude, decay time, and rise time of sEPSC.

### Autoxidation measurement

For measuring the autoxidation of DA and DA-HA, absorbance wavelength at 405 nm was used ^51^. For optimizing the visualization condition, 100 µM DA was added to HEPES, ACSF, and neurobasal media and measured the autoxidation using a microplate reader (GloMax explorer multimode, Promega). For measuring the concentration-dependency of autoxidation without neurons, 1, 5, 10, 50, and 100 µM DA was added and incubated for 24 h and 72 h. For measuring autoxidation with neurons, 1, 5, 10, 50, and 100 µM DA was added and incubated for 7 days. For the experiment for Fig. 6, 200 µM of DA, L-DOPA, 6-OHDA, and DA-HA were added and incubated for 0 to 72 h. At the end of the experiment, absorbance was measured at 405 nm using a microplate reader. All materials were maintained in a humidified atmosphere containing 5 % CO^2^ / 95 % O^2^ at 37°C.

### CCK-8 assay and MTT assay

To assess the effect of DA and DA-HA on cell viability, CCK-8 (DJDB4000X, Dojindo laboratory) or MTT (CellTiter 96® Non-Radioactive Cell Proliferation Assay, Promega) was used. Neurons were seeded on the 96-well plate (8 × 10^4^ neurons / well) with the treatment of 1, 10, 50, and 100 µM of DA and DA-HA for Fig. 5 and 200 µM of DA, L-DOPA, 6-OHDA, and DA-HA for Fig. 6. After 7 days (Fig. 5) and 3 days (Fig. 6) incubation, we added dye solution for 2 h in 5 % CO^2^ / 95 % O^2^ incubator at 37 °C. After that, the absorbance was measured at 450 nm (CCK-8) or 560 nm (MTT) using a microplate reader (GloMax explorer multimode, Promega).

### Immunohistochemistry

The mouse was anesthetized with isoflurane and perfused with 4 % PFA. After perfusion, a whole brain was dehydrated by 30 % sucrose in 0.1 M PBS buffer at 4 °C overnight. After dehydration, 30 µm-thick brain sections were obtained by a cryo-microtome (CM1950, Leica). Sections were firstly incubated with a blocking solution (0.3% Triton-X, 2% of donkey and goat serum in 0.1 M PBS) for 1 h and then immunostained with a primary antibody (chicken anti-TH, ab76442, Abcam) in a blocking solution at 4 °C overnight. The next day, sections were washed with PBS 3 times and incubated with a secondary antibody (Alexa 488 anti-chicken, 703-545-155, Jackson ImmunoResearch) for 1 h at RT and then washed with PBS 3 times for 10 m. DAPI (46190, Pierce) was added to the last washing solution. Sections were carefully transferred onto slide glass and dried briefly. Then fluorescent mounting medium (S3023, Dako) was applied to brain sections and covered by a cover glass. For TH optical density analysis, images were obtained by slide scanner (Axio scan.Z1, Zeiss) and analyzed by ZEN 3.1 blue software (Zeiss). A confocal microscope (LSM 900, Zeiss) was used to obtain images of guide cannula targeting the striatum in Fig 6.

### 6-Hydroxydopamine (6-OHDA) lesions

6-OHDA lesion model was generated as described previously with a little modification ^30^. Briefly, mice were anesthetized by isoflurane (3 % for induction and 1 - 1.5 % during surgery) and were placed in a stereotactic frame (68537, RWD Life Science, Guandong, China). Mice received unilateral injections of 6-OHDA dissolved in 0.02 % ascorbic acid (7.5 μg/ul, 2 μl, 0.2 μl/min) into the left side of the SNpc at the following coordinates: AP, -3.3 mm; ML, -1.3 mm; and DV, -4.2 mm. The body temperature of the mouse was kept at 37 °C during the injection. Guide cannula insertion was performed right after the 6-OHDA injection.

### Guide cannula implantation into striatum

After finishing the 6-OHDA injection, the surface of the skull was dried by an air blower and Optibond (All-In-One, Kerr) was applied throughout the skull. After light curing for 20 seconds, a double guide cannula (C235-3.0/SPC, P1 Technologies) with a dummy cannula (C235DC/SPC, P1 Technologies) was inserted into the striatum using the following coordinates: AP, 0.75 mm; ML, ± 1.25-1.3 mm; and DV, -2.3 mm. After insertion, Charisma (Classic, Kulzer) was applied throughout the skull as well as the guide cannula. After applying UV light for 20 seconds, the skull and guide cannula was tightly fixed together. Finally, dental cement (Self-curing set, Vertex) was applied to complete guide cannula implantation. The mouse was individually housed after surgery. To reduce pain and inflammation, Meloxicam (3 mg/kg, i. p., Metacam, Boehringer Ingelheim) was treated once a day for 3 days including the day of surgery.

### Apomorphine-induced rotation test

One day before the test, the mouse was placed in cylinders (a diameter of 20 cm and height of 25 cm, transparent) which is enclosed by a soundproof chamber. Mouse freely explored inside of cylinder for 10 m for habituation. On Day 1, apomorphine hydrochloride (0.02 mg/kg, 2073, Tocris) was delivered subcutaneously to a mouse and transferred into the cylinder chamber immediately after apomorphine injection. About 5 m after injection, apomorphine-induced rotation behavior was recorded for 30 m. On Day 2, mice were anesthetized with isoflurane (3 % for induction and 0.5 - 1 % during injection) and a dummy cannula was removed. A double internal cannula (C235I/SPC, P1 Technologies) was connected to mineral oil (M8410, Sigma) filled with a syringe (84855, Hamilton) by polyethylene tubing (PE30, PE50, and PE90, BD Intramedic). The syringe was mounted on a micro-infusion pump (Legato 130, KD Scientific) and 0.02 % ascorbic acid was loaded into the syringe. The internal cannula was inserted into the guide cannula and infused 0.02 % ascorbic acid (2 μl, 0.2 μl/min) into the striatum. During infusion, the level of anesthesia should be light enough to perform a rotation test right after infusion of the drug. The mouse was moved into the home cage to recover from anesthesia about 5 m. Then, the rotation test was performed the same as on Day 1. On Day 3, the same behavior procedure as Day 2 was done except that instead of 0.02 % ascorbic acid, 2.5 μM DA-HA, 2.5 μM L-DOPA, or 2.5 μM DA-HA was delivered into the striatum. Lastly, on Day 4, the rotation test was performed to investigate the effect of the drug after 24 h. The number of counterclockwise rotations was measured for full body rotations (360 °).

### Statistical analysis

The numbers and individual dots refer to the number of cells or animals unless otherwise clarified in the figure legends. For data presentation and statistical analysis, GraphPad Prism 9.1.2 (GraphPad Software) was used. For the electrophysiological experiment, Minianalysis (Synaptosoft) and Clampfit (Molecular Devices) were used. For plotting electrophysiological data, Sigma plot (Systat) was used. Statistical significance was set at *p < 0.05, **p < 0.01, ***p < 0.001, ****p < 0.0001. Data are presented as mean ± SEM.

## Acknowledgement

This work is supported by the Intramural Research Program (1711133843) of the Korea Institute of Toxicology (KIT) to S. W. K.; by the Institute for Basic Science (IBS), Center for Cognition and Sociality (IBS-R001-D2) funded by the Ministry of Science to C.J.L.; and by a grant 20182MFDS423 (1475012885) from the Ministry of Food and Drug Safety in 2022, a grant from the Korea Institute of Toxicology (KIT) Research Program (1711159828) and a grant from R&D Convergence Program of the National Research Council of Science and Technology (NST) (CAP-18-02-KRIBB) of Republic of Korea to D. H. W. We thank Dr. Li Yulong for providing original clone of GRAB^DA2m^ sensor.

## Author contributions

D. H. W. and C. J. L. designed and supervised the study. S. K. performed *ex vivo* imaging, *in vivo* immunostaining, and behavior experiments. Y. K. performed rat primary cell culture and various *in vitro* experiments including imaging, cell viability, and electrophysiological experiments. K. H. P., K. M. H., and S. W. K. synthesized DA-HA. S. K. and Y. K. analyzed the data. S. K., Y. K., D. H. W., and C. J. L. wrote the manuscript.

