## Supplemental file for "Dopamine-Modified Hyaluronic Acid (DA-HA) As A Novel Dopamine-Mimetics With Minimal Autoxidation And Cytotoxicity"

[1] KU-KIST Graduate School of Converging Science and Technology, Korea University, Seoul 02841, South Korea, [2] Center for Cognition and Sociality, Cognitive Glioscience Group, Institute for Basic Science, Daejeon 34126, South Korea,[3] Human and Environmental Toxicology, University of Science and Technology, Daejeon 34114, South Korea,[4] Department of Advanced Toxicology Research, Korea Institute of Toxicology, KRICT, Daejeon 34114, South Korea, [5] Department of Polymer Science and Engineering, Chungnam National University, Daejeon 34134, South Korea,[6] Research Group for Biomimetic Advanced Technology, Korea Institute of Toxicology (KIT), KRICT, Daejeon 34114, South Korea

**Keywords:** Dopamine-Modified Hyaluronic Acid (DA-HA), dopamine (DA), DA-induced cytotoxicity, autoxidation, and 6-OHDA-induced mouse model of Parkinson’s disease.

† these are equally contributed

* Correspondence

**To whom correspondence should be addressed:**

**Changjoon Justin Lee, Ph.D.**

Center for Cognition and Sociality, Cognitive Glioscience Group,

Institute for Basic Science, Daejeon 34126, South Korea.

TEL: + 82-42-878-9150, FAX: + 82-42-878-9151

**Dong Ho Woo, Ph.D.**

Department of Advanced Toxicology Research,

Korea Institute of Toxicology, KRICT, Daejeon 34114, South Korea.

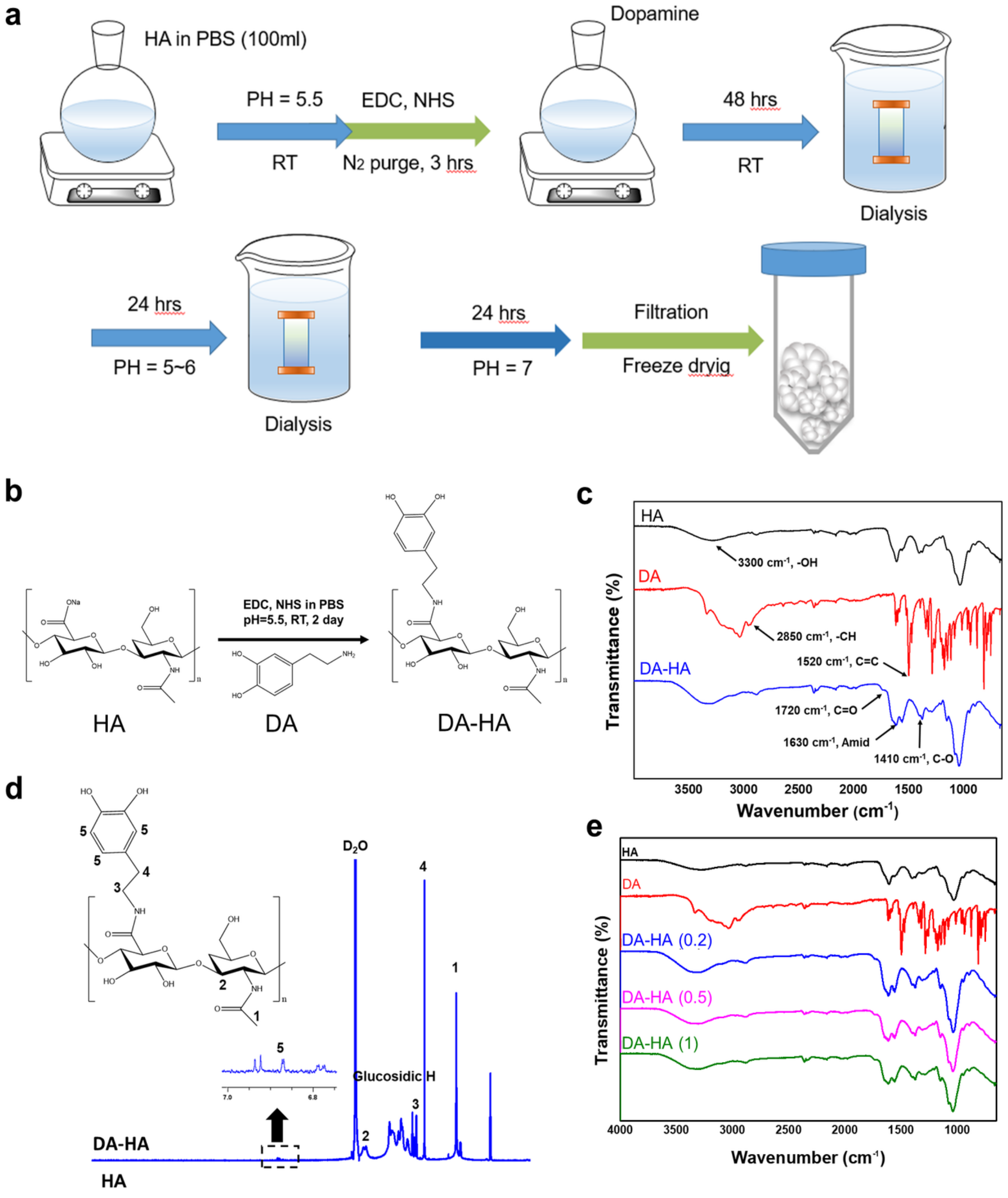


**Fig. S1 The process of synthesis and characterization of DA-HA. a** The process of synthesis of DA-HA. **b** Schematic representation of DA-HA. **c** Protein conformation of HA (black), DA (red), and DA-HA (blue) is analyzed by ATR-FTIR spectra. **d** The structures of HA and DA-HA were analyzed by H-NMR spectra. **e** ATR-FTIR spectra of HA, DA, 0.2 DA-HA, 0.5 DA-HA, and 1 DA-HA.

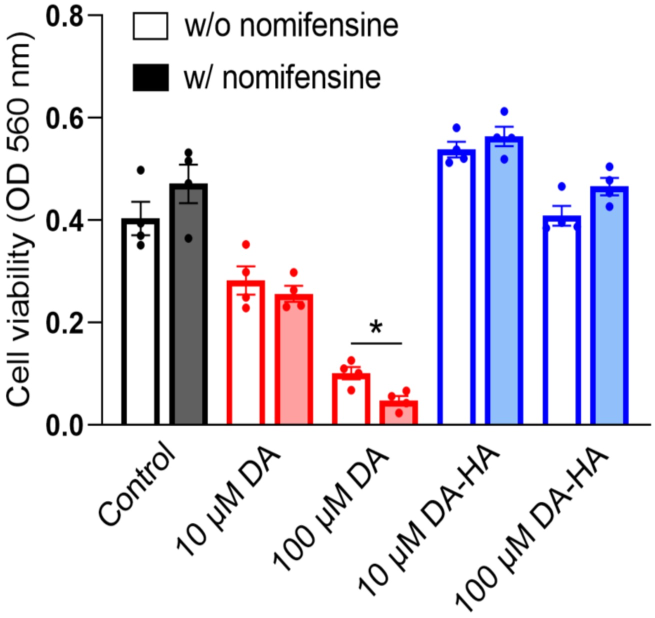


**Fig. S2 Intracellular ROS does not contribute to DA-induced neurotoxicity.** Summary bar graph of cell viability with and without 10 µM nomifensine, a DA transporter (DAT) inhibitor, in the presence of 10 µM DA, 100 µM DA, 10 µM DA-HA, and 100 µM DA-HA for 7 days. At 10 µM DA, nomifensine shows no neurotoxic effect. At 100 µM DA, nomifensine even exacerbates neurotoxicity, indicating that intracellular ROS does not contribute to DA-induced neurotoxicity. Paired t-test. $*p < 0.05$.

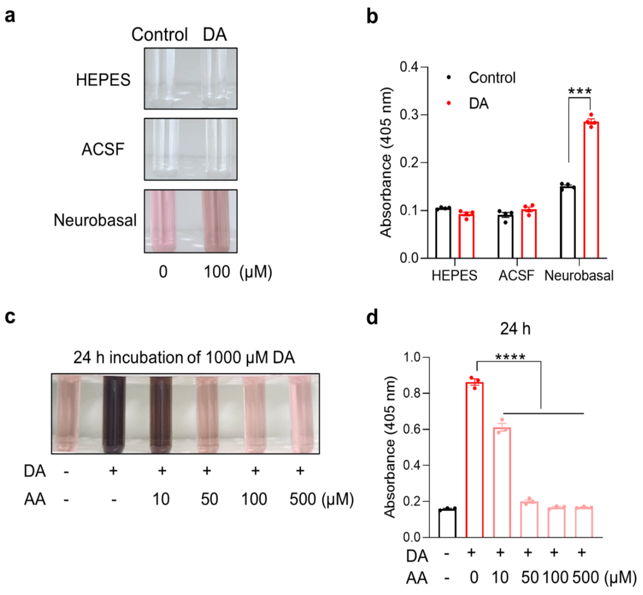


**Fig. S3 Detection of ascorbic acid-sensitive autoxidation in culture media. a** Photopraph of DA autoxidation in various media. Neurobasal media was suitable for visualizing the autoxidation level of 100 µM DA. **b** Summary bar graph of the level of DA autoxidation in HEPES, ACSF buffer, and neurobasal media (n = 4 for each condition). Unpaired t-test. Control vs DA in neurobasal condition, ***$p < 0.001$. **c** Representative photographs of dose-dependency of ascorbic acid (AA) on the autoxidation of 1000 µM DA. **d** Autoxidation of 1000 µM DA with a dose-dependency of AA for 24-h incubation (n = 3 for each condition). One-way ANOVA, $F(5, 12) = 601.6$, $p < 0.0001$; 0 vs 10, 0 vs 50, 0 vs 100, and 0 vs 500 µM AA, ****$p < 0.0001$. Dunnett's multiple comparisons test for **d**.

**Supplementary figures**

| Sample | Feed molar ratio (DA / HA residue) | Degree of conjugated dopamine (%) | Amount of DA per HA (mg / g) | Amount of DA per HA (mmol / g) | Yield (%) |
|---|---|---|---|---|---|
| DA-HA | 0.1 | 1.13 | 5.33 | 0.03 | 59.79 |
| | 0.2 | 5.01 | 23.59 | 0.12 | 60.32 |
| | 0.5 | 7.51 | 35.33 | 0.19 | 59.91 |
| | 1 | 8.29 | 39.02 | 0.21 | 47.61 |

**Table. 1 Result of the feed molar ratio DA and HA from the synthesis of DA-HA.**
